# Gold Nanoprisms as Optical Coherence Tomography Contrast Agents in the Second Near Infrared Window for Enhanced Angiography in Live Animals

**DOI:** 10.1101/322545

**Authors:** Peng Si, Edwin Yuan, Orly Liba, Yonatan Winetraub, Siavash Yousefi, Elliott Daniel SoRelle, Derek William Yecies, Rebecca Dutta, Adam de la Zerda

## Abstract

Optical coherence tomography angiography (OCTA) is an important tool for investigating vascular networks and microcirculation in living tissue. Traditional OCTA detects blood vessels via intravascular dynamic scattering signals derived from the movements of red blood cells (RBCs). However, the low hematocrit and long latency between RBCs in capillaries makes these OCTA signals discontinuous, leading to incomplete mapping of the vascular networks. OCTA imaging of microvascular circulation is particularly challenging in tumors due to the abnormally slow blood flow in angiogenic tumor vessels and strong attenuation of light by tumor tissue. Here we demonstrate *in vivo* that gold nanoprisms (GNPRs) can be used as OCT contrast agents working in the second near infrared window, significantly enhancing the dynamic scattering signals in microvessels and improving the sensitivity of OCTA in skin tissue and melanoma tumors in live mice. This is the first demonstration that nanoparticle-based OCT contrast agent work *in vivo* in the second near infrared window, which allows deeper imaging depth by OCT. With GNPRs as contrast agents, the post-injection OCT angiograms showed 41% and 59% more microvasculature than pre-injection angiograms in healthy mouse skin and melanoma tumors, respectively. By enabling better characterization of microvascular circulation *in vivo*, GNPR-enhanced OCTA could lead to better understanding of vascular functions during pathological conditions, more accurate measurements of therapeutic response, and improved patient prognoses.

---

Real-time imaging of vascular networks and microcirculation in living tissue plays a crucial role in better understanding pathological conditions.^1^ In particular, better characterization of tumor vasculature *in vivo* could provide significant prognostic value for cancer patients^2^ and improved measurements of their therapeutic responses,^3–6^ since tumor vessels are critical sites for drug delivery, angiogenic therapy, chemotherapy, and immunotherapy.^7^ The variety of imaging tools currently available for *in vivo* angiography can mostly be classified into two categories: macroscopic and microscopic. Macroscopic imaging tools, such as computed tomography, positron emission tomography, ultrasound imaging, and magnetic resonance imaging, provide full body penetration but lack the spatial resolution to resolve small blood vessels in tissue.^8^ Microscopic imaging modalities, such as confocal and multiphoton microscopy, provide subcellular imaging resolution but have suboptimal tissue penetration depth and small fields of view, which limit their applications in clinical settings.^9^ The dilemma presented by these shortcomings of microscopic and macroscopic methods calls out for the development of imaging modalities that can bridge the gap and combine high resolution with deeper penetration.

Optical coherence tomography angiography (OCTA) has emerged in recent years as one of the most powerful and clinically indispensable imaging tools thanks to its subcellular resolution, millimeter tissue penetration depth, and three-dimensional imaging capability. Traditional OCTA detects blood vessels by computing an induced optical signal variance^10, 11^ of moving red blood cells (RBCs), such as the speckle intensity variance, phase variance, or complex field variance.^11^ However, OCTA based on detecting these dynamic scattering signals of RBCs has several intrinsic drawbacks that compromise its sensitivity and limit its utility in skin and tumors. First, the RBC flow in capillaries is in single-file and is highly heterogeneous over time and space.^12–15^ At diverging bifurcations, the capillaries with lower flow rate tend to have lower hematocrit,^16^ which results in long latency between RBCs and discontinuous OCTA signals.^17^ Increasing the OCT B-scan repetitions can improve the OCTA sensitivity for capillaries, but significantly lengthens the image acquisition time. In some capillaries, no RBCs are present due to the ultraslow plasma flow. These capillaries are called plasma channels,^18^ which cannot be detected by traditional OCTA due to the lack of dynamic scattering signals. In addition, RBCs tend to migrate away from vessel walls, leaving a cell-free layer (CFL) in microvessels.^18^ This prevents OCTA from being used as a quantitative tool to accurately measure the vessel diameters. In tumor tissue, strong light attenuation makes it even more challenging to image the microvessels located in the central tumor region. To circumvent these limitations and achieve ultrahigh-sensitivity OCTA for skin and tumor tissues, it is highly compelling to introduce intravascular contrast agents with suitable light-scattering, hydrodynamic, and pharmacokinetic properties.^16^

Gold nanoparticles are attractive candidates as OCT contrast agents due to their high refractive index sensitivity, biocompatibility, and tunable plasmonic resonance.^19^ We recently reported that large gold nanorods can be used as OCT contrast agents in the first near-infrared (NIR-I) window (800–1000 nm).^20^ However, very few gold nanostructures have been reported as OCT contrast agents in the second near-infrared (NIR-II) imaging window (1100–1400 nm), which has lower tissue scattering and deeper photon penetration.^21–24^ In the work reported here, we have demonstrated that gold nanoprisms (GNPRs) are strong scatterers of light in the NIR-II window, and employed them as OCT contrast agents in live animals for the first time. Due to their surface plasmon resonance and high scattering coefficient in the NIR-II window,^25^ GNPRs significantly enhanced the dynamic scattering signals in blood vessels and drastically improved the sensitivity of OCTA for skin tissue and melanoma tumors. After administering GNPRs intravenously, 41% and 59% more microvessels were detected by OCTA in healthy mouse ear skin and melanoma tumors, respectively. No leakage of the nanoparticle-contrast agents were detected outside the blood vessels, suggesting the contrast agent-enhanced OCTA processes high specificity.

GNPRs were prepared according to the method described in Supporting Information (SI). Electron micrographs (Figure S1a,b) shows that the GNPRs have triangular platelet morphology, with an edge length of ~140 nm and thickness of 8 nm. Dynamic light scattering (DLS) measurements (Figure S1c) suggests the GNPRs have uniform size distribution and high yield purity. The surface plasmon resonance peak of the GNPRs was 1385 nm with a full width at half maximum (FWHM) of 375 nm (Figure S1d), which falls in the spectral window of our long-wavelength OCT system (1250–1400 nm). We PEGylated the GNPRs for use in all of our OCT experiments. The OCT scattering signal of individual GNPRs can be visualized at low concentrations (50 and 500 fM) in water in capillary tube phantoms (Figures 1a and S2a). The data was fitted well to a physical model we reported previously^26^ (green curve in Figure 1c, R^2^=0.999), showing that the mean OCT amplitude of the GNPR solution increased linearly with its concentration from 0 to 5 pM (blue dashed line in Figure 1c, R^2^=0.982), and in proportion to the square root of its concentration ranging from 50 pM to 500 pM (red dashed line in Figure 1c, R^2^=0.975). For GNPRs in mouse whole blood samples (Figures 1b and S2b), the highly heterogeneous scattering signals of blood prevented the OCT signals of individual GNPRs from being detected at low concentration (i.e., 100 pM). However, strong OCT signals of GNPRs were imaged in whole blood at concentrations ≥250 pM. We showed that the OCT signal of GNPRs in blood increased with concentration, and the experimental data matched well with the prediction of our physical model^26^ (Figure 1d). We calculated the detection limit of GNPRs in blood to be 206 pM at a signal-to-noise ratio of 3. The speckle variance and normalized speckle variance (NSV) signals in water and blood also grew with increasing GNPR concentrations (Figure S3). The NSV signal is calculated by dividing the raw speckle variance by the OCT amplitude in each voxel, which compensates for attenuation of light and facilitates the imaging of speckle variance signals deeper in tissue.

**Figure 1.**
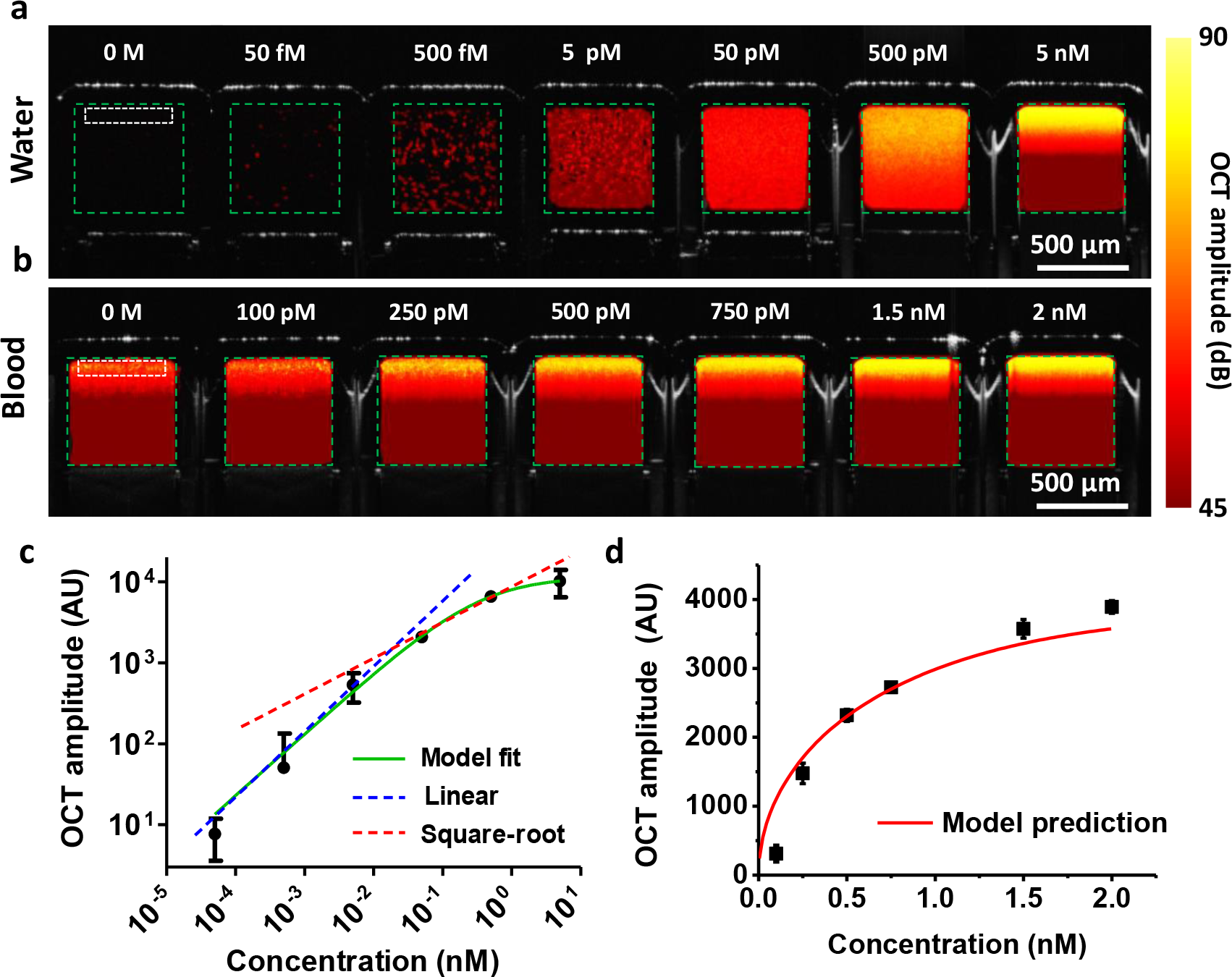
*In vitro* characterization of OCT signals of GNPRs. The OCT signals of different concentrations of PEGylated GNPRs in (a) water and (b) blood in capillary tube phantoms, color-coded by OCT amplitude. The green dashed lines show the inner surface of the square tubes. The white boxes show the regions of interest (ROIs) used to measure the mean OCT amplitude at each GNPR concentration. (c) The mean OCT amplitude as a function of GNPR concentration in water (n = 3 per concentration). Green curve is a fit to a physical model we reported recently.^26^ The blue and red dashed lines indicate the linear and square root regimes of the model, which consider less than one scatterer and multiple scatterers per voxel, respectively. (d) The mean OCT amplitude of GNPRs in blood as a function of concentration (n = 3 per concentration), with the model prediction (red curve). Data are presented as mean ± SEM.

In the first of our *in vivo* experiments we incrementally injected GNPRs into the mouse tail vein (8 injections, each increasing the bloodstream GNPR concentration by 0.25 nM), and acquired B-scan images of the mouse ear (Figure 2a-b) after each injection (Figure S4). (Here and below, all injection concentrations are reported in terms of their calculated increment to the bloodstream GNPR concentration.) To better visualize the OCT signal changes in blood vessels, the vessels were color-coded by OCT amplitude (Figure 2c-k). The OCT signals of blood vessels in the mouse ear increased with increasing GNPR concentration in blood, consistent with the *in vitro* phantom experiment results. We quantitatively measured the OCT signals of 3 blood vessels (regions denoted by dashed circles in Figure 2c) with different GNPR concentrations, showing that the blood vessels with only 0.25 nM GNPRs exhibited significantly greater OCT intensity than they did before injection (Figure 2l, *P* = 0.032). However, the increase of OCT amplitude with each injection was not significant for GNPR concentrations in blood ≥0.75 nM, which can be explained by a multi-scattering effect according to our physical model.^26^

**Figure 2.**
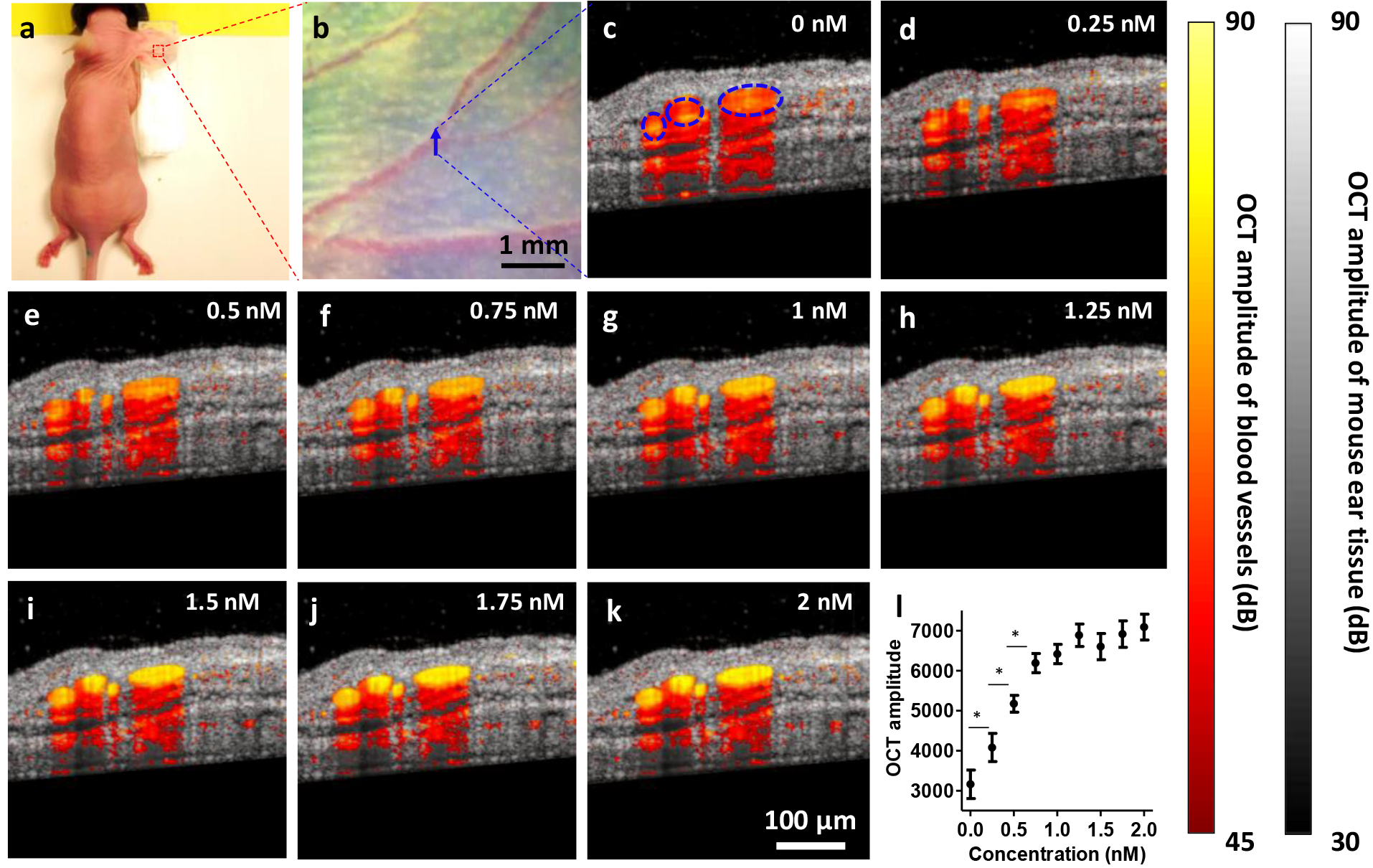
*In vivo* characterization of OCT signals of GNPRs in blood vessels. (a-b) Photographs of imaged live mouse and its pinna. The location of OCT cross-section scans is denoted by arrow in (b). (c-k) Cross-sectional OCT images of the mouse pinna before and after incremental intravenous injections of PEGylated GNPRs. The blood vessels were detected by speckle variance. The OCT signals in the blood vessels are color-coded by OCT intensity. Dashed circles in (c) show the ROIs used to quantify the OCT signals. (l) The OCT amplitude in the blood vessels at various GNPR concentrations (n = 3 vessels). Data are presented as the mean ± SEM, * *P* < 0.05.

Next we investigated OCT angiograms of healthy mouse ears (Figure 3a-c) with increasing concentrations of GNPRs. We obtained the OCT angiograms by detecting the blood vessels with NSV followed by segmenting the vessels with a hybrid intensity/shape-based compounding algorithm.^27^ We show both the en-face projection (Figure 3d1-l1) and cross-section OCT angiograms (Figure 3d2-l2) of a healthy mouse ear before and after incremental injections of GNPRs. With increasing GNPR concentrations in the blood, a higher density of blood vessels can be observed on the OCT angiograms (Figure 3d-l). The volumetric angiogram vessel density (VAVD) was calculated by dividing the total number of vessel voxels of each 3D OCT angiogram by the total number of tissue voxels (Figure 3m). The VAVD of the healthy ears dramatically increased by 33% when the GNPR concentration in blood increased from 0 to 0.75 nM. Statistical analysis shows that the first 0.25 nM GNPRs introduced into the blood increased the VAVD significantly (*P* < 0.05).

**Figure 3.**
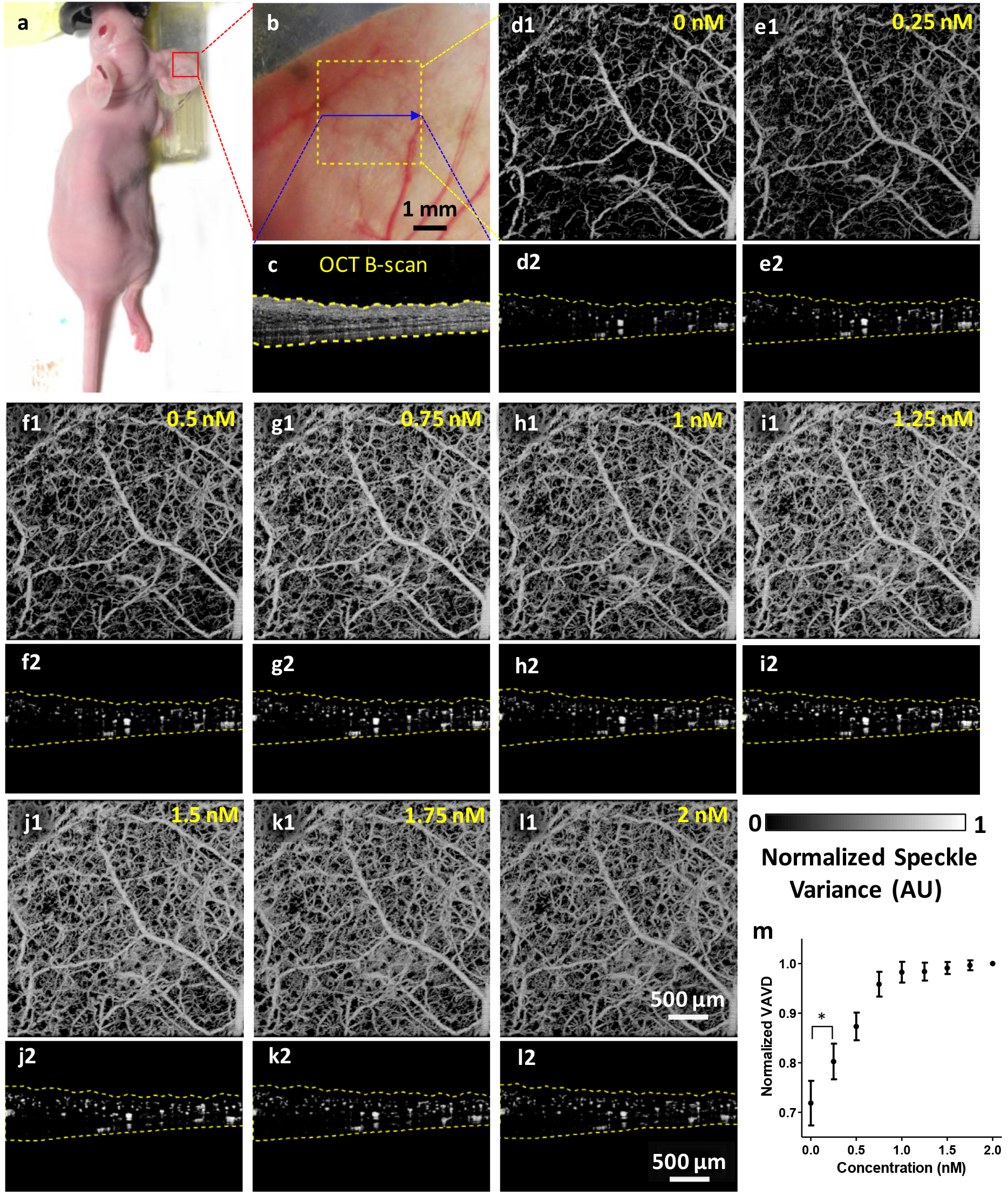
OCT angiograms of a mouse ear before and after incremental injections of GNPRs. (a-b) Photographs of imaged live mouse and magnified mouse pinna. The arrow and box in (b) show the location of OCT cross-section scans and field of view of OCT angiograms, respectively. (c) OCT B-scan image of the tissue microstructure at arrow-denoted location. The yellow dashed lines indicate the tissue boundaries. (d1-l1) En-face projected OCT angiograms of the mouse ear with incremental GNPR concentrations in blood. (d2-l2) Cross-section OCT angiograms of the mouse ear with incremental GNPR concentrations in blood. (s) Normalized volumetric angiogram vessel densities (VAVDs) calculated from 3D OCT angiograms and 3D OCT tissue microstructure images at different GNPR concentrations (n = 3 mice). Data are presented as mean ± SEM, * *P* < 0.05.

We further investigated blood circulation time of GNPRs by obtaining OCT angiograms before injection and at different time points after injection of 0.75 nM GNPRs (Figure S5a-e). The quantification shows that the VAVD peaked at 5 min post-injection, and gradually decreased over time until 24 h post-injection (Figure S5f). The result suggests that GNPRs were washed out from the mouse pinna blood vessels in 24 h.

We next investigated if the sensitivity of OCTA can be influenced by bolus injection of vehicle or the use of different OCTA algorithms. We sequentially injected 100 μL vehicle (water) and 100 μL of GNPRs in mice via tail veins, and took 3D OCT images of the mouse ears before injection and 5 min after each injection. The corresponding OCT angiograms were computed by both the speckle variance–based OCTA algorithms (Figure 4a-c) and complex variance–based optical microangiography (OMAG)^28^ (Figure S6), which is more sensitive to slow blood flows that have only phase changes.^16^ After vehicle injection, we observed a slight reduction of vessel density on the OCT angiograms computed by both algorithms (Figures 4b and S6b), possibly because the increased blood plasma volume slightly reduced the hematocrit in the capillaries. Significantly more vascular plexuses can be seen on the OCT angiograms after injecting the GNPRs (Figures 4c and S6c). The arrows in the magnified images (Figure 4h,i) indicate capillaries that were not observed on the OCT angiograms before and after vehicle injection (Figure 4d-g), but clearly present on the OCT angiogram after injecting GNPRs. Quantitative analysis shows that the VAVD decreased by 8% after vehicle injection but increased by 45% following the GNPR injection (Figure 4m). No significant vessel density differences were observed between speckle variance– and complex variance–based angiograms, indicating that different OCTA algorithms have negligible influence on the OCTA sensitivity compared to the injection of contrast agents.

**Figure 4.**
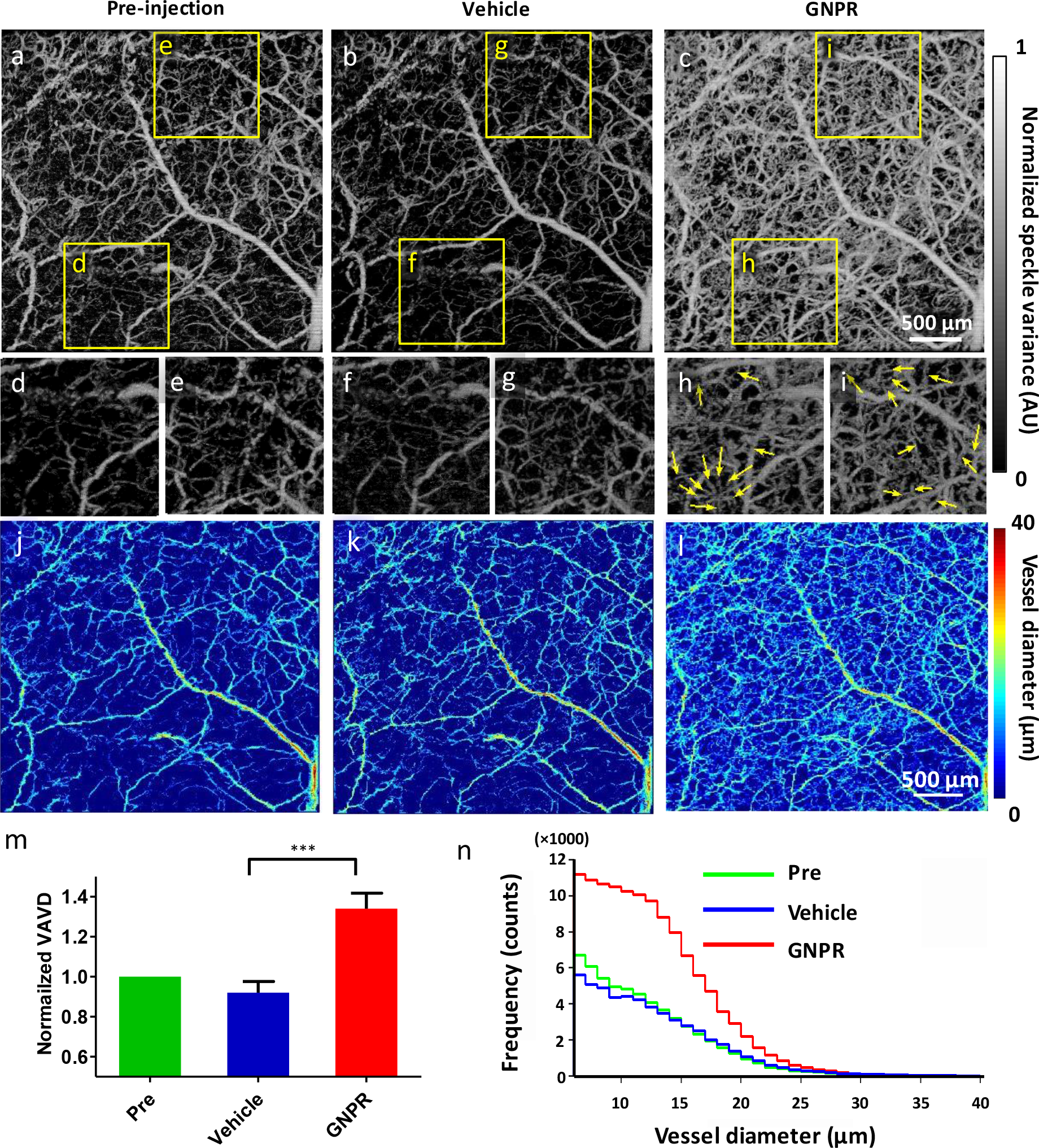
OCT angiograms and vessel diameter maps of a mouse ear before injection, after vehicle injection, and after 0.75 nM GNPR injection. (a-c) En-face projected OCT angiograms computed by speckle variance–based algorithms of the mouse ear (a) before injection, (b) after vehicle injection, and (c) after subsequent GNPR injection. See Figure S6 for corresponding images of the same region computed by complex variance–based OCTA. (d-i) Magnified microvasculature networks in the boxes of (a-c). The yellow arrows in (h,i) denote capillaries that were observed only after GNPR injection. (j-l) Vessel diameter maps corresponding to the OCT angiograms. (m) Normalized VAVDs. n = 3 mice, *** *P* < 0.001. (n) Histogram of the vessel diameter distribution in the OCT angiograms.

Comparable NSV signals were observed in blood vessels before and after vehicle injection, but significantly higher NSV signals can be observed in the vessels after GNPR injection (Figure S7a-g). The cross-section angiograms showed similar vessel quantity and comparable vessel sizes before and after vehicle injection, but a significantly higher number of capillaries (diameter <10 μm) and larger-diameter vessels can be observed after GNPR injection (Figure S7h-n). This improvement can be attributed to the strong dynamic scattering signals of GNPRs and their homogeneous spatial distribution across the microvascular networks, which dramatically improved the OCT sensitivity and allowed the imaging of capillaries with ultraslow flow, ultralow hematocrit, plasma channels, and CFLs. The vessel diameter maps (Figure 4j-l) and histogram (Figure 4n) showed that the injection of vehicle slightly reduced the number of capillaries visible in the OCT angiograms without altering the number of larger-diameter vessels visible. This reduced OCTA sensitivity could be caused by decreased hematocrit in capillaries due to dilution. In contrast, after GNPR injection the angiograms of mouse ears showed significantly more capillaries, indicating GNPR-enhanced OCTA dramatically improved the detection of capillaries with low RBC dynamic scattering signals, such as plasma channels and capillaries with low hematocrit and ultraslow blood flow. We also noticed that the GNPR-enhanced OCTA detected more non-capillary microvessels. This might occur because the enhanced OCTA detects CFLs, improving the detection of vessels and increasing their measured diameters. By providing strong dynamic scattering signals throughout the entire vascular network, intravascular GNPRs lead to significantly enhanced sensitivity of OCTA.

To further investigate if the OCTA signals were predominantly derived from GNPR-induced dynamic scattering, we injected vehicle and GNPRs, respectively, in two groups of mice and compared their OCT angiograms before and after sacrificing the mice (Figure S8). The post-sacrifice images were obtained 30 min after the mice’s hearts stopped beating. For the control group, the microvascular plexuses were clearly visible on the OCT angiograms before sacrifice, both before and after vehicle injection (Figure S8a,b). However, only weak OCTA signals of the largest blood vessels were seen on the angiograms obtained after the mice’s deaths (Figure S8c) due to the cessation of RBC flow. For the GNPR-injected mice, the OCT angiograms showed significantly more OCTA signals of microvascular plexuses (Figure S8d,e), which barely decreased at 30 min after the mice’s hearts stopped beating (Figure S8f). This result suggests the OCTA signals of the GNPR-injected mice mainly originated from the intravascular dynamic scattering of GNPRs, rather than RBCs. The absence of RBC dynamic motion has negligible influence on the GNPR-enhanced OCTA signals.

The imaging specificity of GNPR-enhanced OCTA was investigated with two-photon microscopy. The mice were intravenously injected with dextran-rhodamine followed by Alexa488-labeled GNPRs. Two-photon microscopy images of the mouse ear taken 5 min post-injection show no fluorescent signals of Alexa488-labeled GNPRs can be observed outside the blood vessels (Figure S9), indicating there was no extravasation of the contrast agents and GNPR-enhanced OCTA has high specificity for imaging the microvasculature network.

To investigate how GNPR-enhanced OCTA can improve the microvascular imaging in tumors, an orthotopic B16 melanoma mouse model was used for the study (inset of Figure 5a). The mouse ear implanted with melanoma tumors was imaged before and 5 min after injection of 0.75 nM GNPRs (Figure S10). The OCT angiograms were color-coded by depth (Figure 5a-d, Movies S1 and S2). Significantly higher melanoma tumor vasculature is observed on the post-injection OCT angiogram (Figure 5b) than the pre-injection OCT angiogram (Fig.5a, Movie S3). From the cross-section view, the GNPR-enhanced OCT angiogram not only shows markedly more microvasculature at the tumor-host interface (Figure 5c,d), but also shows deeper tumor vessels in the central tumor area (arrows in Fig. 5d), which is 600~800 μm below the skin surface. The post-injection angiogram shows more vessels at the tumor-host interfaces than in the central tumor regions, as is expected since vascular density tends to decrease as tumors grow, leading to ischemia zones inside the core tumor.^29^ More accurate mapping of the tumor vascular network enables better assessment of the ischemia region, which is crucial for evaluating the response to anti-angiogenic therapy. Increased vasculature was also observed in the peritumoral tissue in the post-injection OCT angiogram. We analyzed the vascular density of the tumor and peritumoral tissue at different depths (Figure 5e). For peritumoral tissue, the highest densities of vessels are at 170 and 260 μm beneath skin surface, which locate in the two dermis layers on the dorsal and ventral sides of the ear, respectively. The post-injection angiogram shows ~30% higher vessel densities at each of these two depths. For the melanoma tumoral tissue, 65% more vasculature are shown on the post-injection OCT angiogram compared to the pre-injection angiogram at the depth of 200 μm, which has the highest tumor vascular density. The VAVDs of peritumoral and tumor tissue increased by 28 % and 59 % respectively following the administration of GNPRs (Figure 5f). The VAVD of the tumors was greater than that of the peritumoral tissue, which could possibly be attributed to tumor-induced angiogenesis. The higher percentage increase of tumor VAVD than peritumoral VAVD after contrast agent injection can be explained by the fact that angiogenic tumor vessels having significantly lower blood flow than the normal vessels.^29^

**Figure 5.**
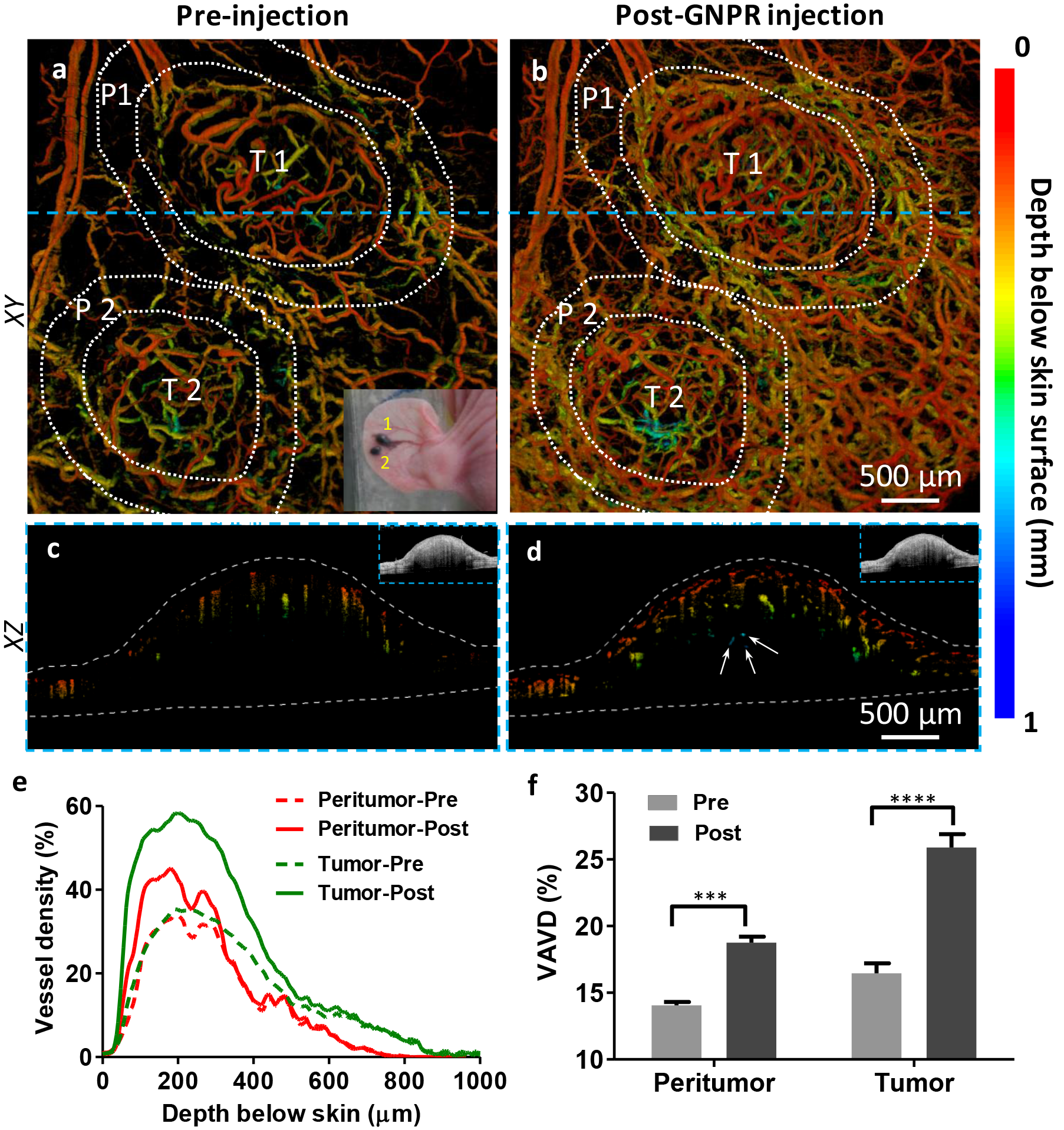
Contrast-enhanced OCT angiography of melanoma tumors. (a,b) En-face projection OCT angiograms (color-coded by depth below skin surface) of melanoma-bearing mouse ear tissue (a) before and (b) 5 min after 0.75 nM GNPR intravenous administration. The tumors and peritumoral regions are circled by dashed lines. T: tumor, P: peritumoral tissue. The inset in (a) shows a photograph of the tumor-bearing ear. The horizontal dashed blue line indicates the B-scan position in (c) and (d). (c,d) Depth color-coded cross-sectional OCT angiograms showing the melanoma vasculature in tumor 1 (c) before and (d) after GNPR injection. The arrows indicate deep tumor vessels detected by OCTA only after GNPR injection. Dotted lines indicate the dorsal and ventral skin surfaces. The insets show B-scan images of the tissue at the same transverse location. (e) The OCT angiogram vessel densities of peritumoral and tumor tissues at different depths beneath the dorsal skin surface before and after GNPR administration. (f) The total VAVDs of peritumoral and tumor tissue before and after GNPR injection (n = 2 tumors), *** *P* < 0.001, **** *P* < 0.0001.

In summary, we have shown that OCTA signals derived from dynamic scattering of RBCs alone cannot accurately depict the complete microvasculature network in skin and tumors. This traditional OCTA is compromised by the high spatial and temporal heterogeneity of RBC flow in small blood vessels (i.e., the presence of plasma channels, low hematocrit, and long latency between RBCs in capillaries). It is particularly challenging to image tumor microvasculatures not only because of the significantly lower overall blood flow in angiogenic tumor vessels compared to normal vessels, but also because of the low scattering signals from the deep tumor vessels due to strong light attenuation. We have demonstrated that the sensitivity of OCTA can be dramatically improved by injecting PEGylated GNPRs as OCT contrast agents in the NIR-II imaging window, which circumvents the limitations of traditional OCT relying on RBC scattering signals and enables better characterization of the vascular networks of skin and tumors.

The surface plasmon resonance of GNPRs synthesized in this work is 1385 nm, which will also allow them to be used as spectral contrast agents^20^ in other applications in the future. However, although GNPR-enhanced OCTA significantly improved microvascular imaging sensitivity in tumors, this technique may not image nonfunctional tumor vasculatures which are not perfused continuously. In addition, OCT’s inherently limited tissue penetration depth might prevent even the GNPR-enhanced OCTA from detecting microvessels too deep in a very large tumor (e.g. ≥ 1 mm in depth).

Nevertheless, the GNPR-enhanced OCTA technique provides a useful tool that can help biological and clinical investigators better characterize the vascular structure and density in skin and tumors *in vivo*. Accurate mapping of vasculature during pathological conditions is crucial for disease progression analysis, therapeutic monitoring, and patient prognoses. Applied to tumors, the technique can help advance understanding of features of the tumor microenvironment such as ischemia, which is an important indicator for tumor progression and response to anti-angiogenic therapy. Furthermore, the concentration-dependent scattering signals provided by GNPRs in the bloodstream will allow GNPR-enhanced OCTA to be used as a quantitative tool for more accurate measurement of plasma flow, transit time, vessel diameters, and so on. These quantitative assessments of blood vessels would enable OCTA to better study pathophysiology and expand its clinical applications.

## SUPPORTING INFORMATION

Additional description of the materials, methods, and results; Figures S1–S10 (PDF)

Movie S1 showing standard OCTA of melanoma tumor, color-coded by depth (AVI)

Movie S2 showing GNPR-enhanced OCTA of melanoma tumor, color-coded by depth (AVI)

Movie S3 showing OCTA of melanoma tumor, color-coded before and after GNPR enhancement (AVI)

## AUTHOR INFORMATION

### Corresponding Author

*

### Author Contributions

The manuscript was written through contributions of all authors. All authors have given approval to the final version of the manuscript.

## ACKNOWLEDGEMENTS

This work was funded in part by grants from the United States Air Force (FA9550-15-1-0007), the National Institutes of Health (NIH DP50D012179), the National Science Foundation (NSF 1438340), the Damon Runyon Cancer Research Foundation (DFS# 06-13), Claire Giannini Fund, the Susan G. Komen Breast Cancer Foundation (SAB15-00003), the Mary Kay Foundation (017-14), the Skippy Frank Foundation, the Donald E. and Delia B. Baxter Foundation, a seed grant from the Center for Cancer Nanotechnology Excellence and Translation (CCNE-T; NIH-NCI U54CA151459), and a Stanford Bio-X Interdisciplinary Initiative Seed Grant (IIP6-43). We would like to thank Stanford Neuroscience Microscopy Service (NMS), supported by NIH NS069375 and Stanford Cell Sciences Imaging Facility (CSIF).

